# Targeting the selectivity filter to drastically alter the activity and substrate spectrum of a promiscuous metal transporter

**DOI:** 10.1101/2025.05.07.652779

**Authors:** Yuhan Jiang, Michael Nikolovski, Tianqi Wang, Keith MacRenaris, Thomas V. O’Halloran, Jian Hu

**Author notes:** Corresponding authors: Thomas V. O’Halloran; Jian Hu. Equally contributive to this work (Yuhan Jiang, Michael Nikolovski).

## Abstract

*d*-Block metal transporters play a crucial role in maintaining the homeostasis of life-essential trace elements and are attractive targets for protein engineering aimed to selectively enriching or excluding metals in living organisms. However, systematic efforts to engineer these transporters have been hindered by limited understanding of their transport mechanism and substrate specificity. In this study, we applied a focused-screen approach to human ZIP8, a promiscuous *d*-block divalent metal transporter, by systematically changing three key residues that form the selectivity filter at the entrance of the transport pathway. Screening a library of 48 constructs using an ICP-MS-based transport assay, we identified variants with significantly altered transport activities and/or substrate preferences. The E343D variant exhibited dramatically enhanced activity for all tested metal substrates, a shift in substrate preference, and an expanded substrate spectrum including the non-substrate metals VO^2+^ and Cu^2+^. Additionally, we identified lead ion (Pb^2+^) as a substrate of wild-type ZIP8. These findings suggest that the ZIP fold is highly adaptable and amenable for transporting a wide range of metals with diverse physicochemical properties, making it a promising scaffold to generate novel metal transporters for applications.

## Introduction

The *d*-block metals manganese, iron, cobalt, nickel, copper, zinc, and molybdenum perform essential functions in biological systems by playing structural, catalytic, and regulatory roles in various biomolecules.^1-7^ The numerous physiological roles of these metals and the potential deleterious effects upon dysregulation underscore the need for precise control over their concentrations and distributions within living organisms. Metal transporters, which function as selective gates to regulate the flow of metal ions across biological membranes, are central players in maintaining the homeostasis of these micronutrients at cellular and systemic levels.^8, 9^

The divalent *d*-block metal transporters are considered to be attractive engineering targets for applications such as biofortification,^10, 11^ phytoremediation,^12, 13^ biomining,^14, 15^ and heavy metal exclusion from food.^16, 17^ From this perspective, the Zrt-/Irt-like protein (ZIP) family has attracted increasing attention due to its ubiquitous expression in all kingdoms of life, its broad substrate spectrum and, in particular, its pivotal role in metal uptake from the environment.^18-22^ Creating ZIPs with desired properties will allow a highly selective accumulation or exclusion of specific metals in the host organism. For example, engineering IRT1, a plant root-expressing iron-transporting ZIP, to eliminate Cd transport activity while retaining iron transport activity may help reduce Cd uptake by crops grown on the contaminated lands.^23^

In our previous study of ZIP8, a promiscuous ZIP that transports Zn^2+^, Fe^2+^, Mn^2+^, Co^2+^, and Cd^2+^,^24-30^ we showed that a combination of four mutations led to a variant with increased Zn^2+^ transport activity and drastically reduced activities for Cd^2+^, Fe^2+^, and Mn^2+^, demonstrating the feasibility of rational engineering of a ZIP transporter.^31^ The strong epistatic interaction between the two residues selected for mutagenesis allowed us to identify a selectivity filter located at the entrance of the transport pathway (**Figure 1**). E318 and E343 from the transport domain approach Q180 in the scaffold domain when the transporter adopts an outward-facing conformation, according to the proposed elevator transport mode,^32-34^ and form a transient metal binding site that screens incoming metal ions before reaching the transport site. Bioinformatics analysis of these three positions showed that, although the selectivity filter-like structure is likely present in many ZIPs, the amino acid composition is highly variable,^31, 35^ suggesting that these transporters may have different substrate preferences, and that the selectivity filter represents an ideal target to alter the substrate spectra of ZIPs.

**Figure 1.**
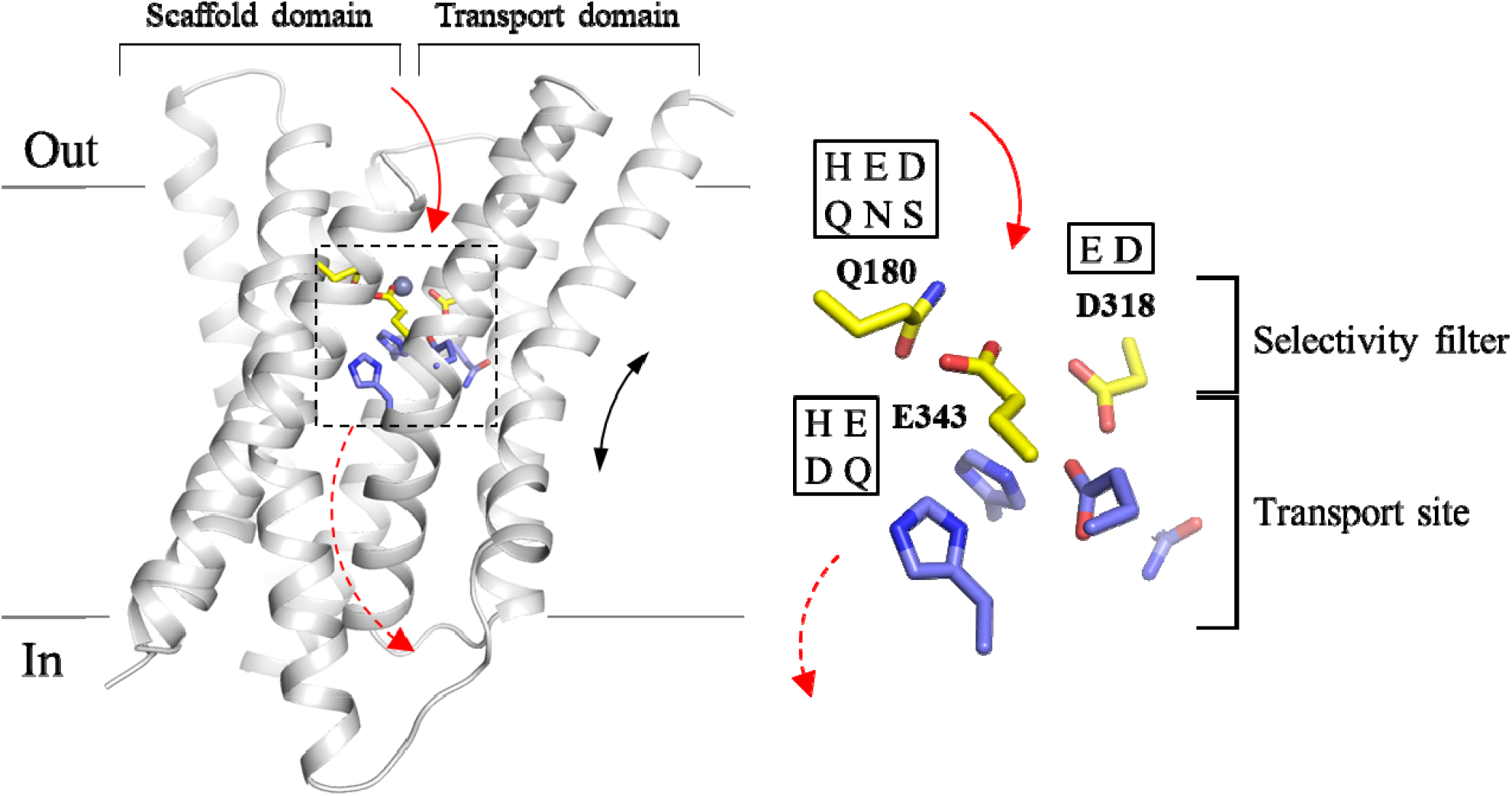
The selectivity filter and the transport site of ZIP8. *Left*: the structural model of human ZIP8 in the outward-facing conformation. The solid arrow indicates the pathway for metal to enter the transport site through the selectivity filter, and the dashed arrow indicates the pathway for metal to be released to the cytoplasm when the transporter switches to the inward-facing conformation. The black double-headed arrow indicates the elevator motion of the transport domain relative to the scaffold domain during transport. The model was generated by homology modeling using the AlphaFold predicted ZIP13 structure (in an outward-facing conformation) as template (UniProt ID: Q96H72). For clarity, long loops connecting transmembrane helices are trimmed. The residues at the selectivity filter are shown in yellow and the residues forming the transport site are in light blue. The metal substrate is depicted as a grey sphere. D318 and E343 contribute to both the selectivity filter and the transport site. *Right*: the zoomed-in view of the selectivity filter and the transport site. The one-letter codes of the amino acids tested at the indicated positions in this work are shown in the frames.

In this work, we explored the extent to which the transport properties of ZIP8 can be tuned by systematically altering the amino acid composition of the selectivity filter and screening the variants against a panel of *d*-block metals using an ICP-MS-based transport assay.^36^ This approach allowed us to identify the variants with drastically altered activity and substrate preference, demonstrating the great potential of ZIPs as targets of transporter engineering for selective metal transport in applications.

## Results

### Construction of a ZIP8 variant library

Since the selectivity filter is at the forefront of metal interaction, we hypothesized that changing the amino acid composition of the selectivity filter would significantly change substrate preference. To test this, we systematically introduced polar amino acid residues at these positions based on the bioinformatics analysis of the ZIP family (**Figure 1**).^35^ Q180 is at the pore entrance and this position is mostly occupied by a polar or charged residue, including histidine, aspartate, glutamate, asparagine, glutamine, and serine.^31^ E318 is one of the residues at the transport site and part of the selectivity filter. In other ZIPs, this position can also be occupied by aspartate or histidine.^35^ A histidine at this position is present only in ZIP13, an ER/Golgi-residing ZIP that was shown to be a zinc importer^37^ but then identified as an iron exporter.^38-40^ In this work, histidine was not introduced to this position because the selectivity filter-like structure in ZIP13 is unlikely to screen metals if ZIP13 maintains the same topology as other ZIPs but functions as an exporter. E343 is another residue playing roles in both the selectivity filter and the transport site. In addition to glutamate, the position of E343 can also be occupied by histidine or glutamine in other ZIPs^31^. Given the similar property to glutamate, aspartate was also introduced to this position although it is not present at this position in any known ZIP. Eventually, a library consisting of 48 constructs (wild-type ZIP8 and 47 variants) was generated for functional screen (**Table 1**).

**Table 1.**
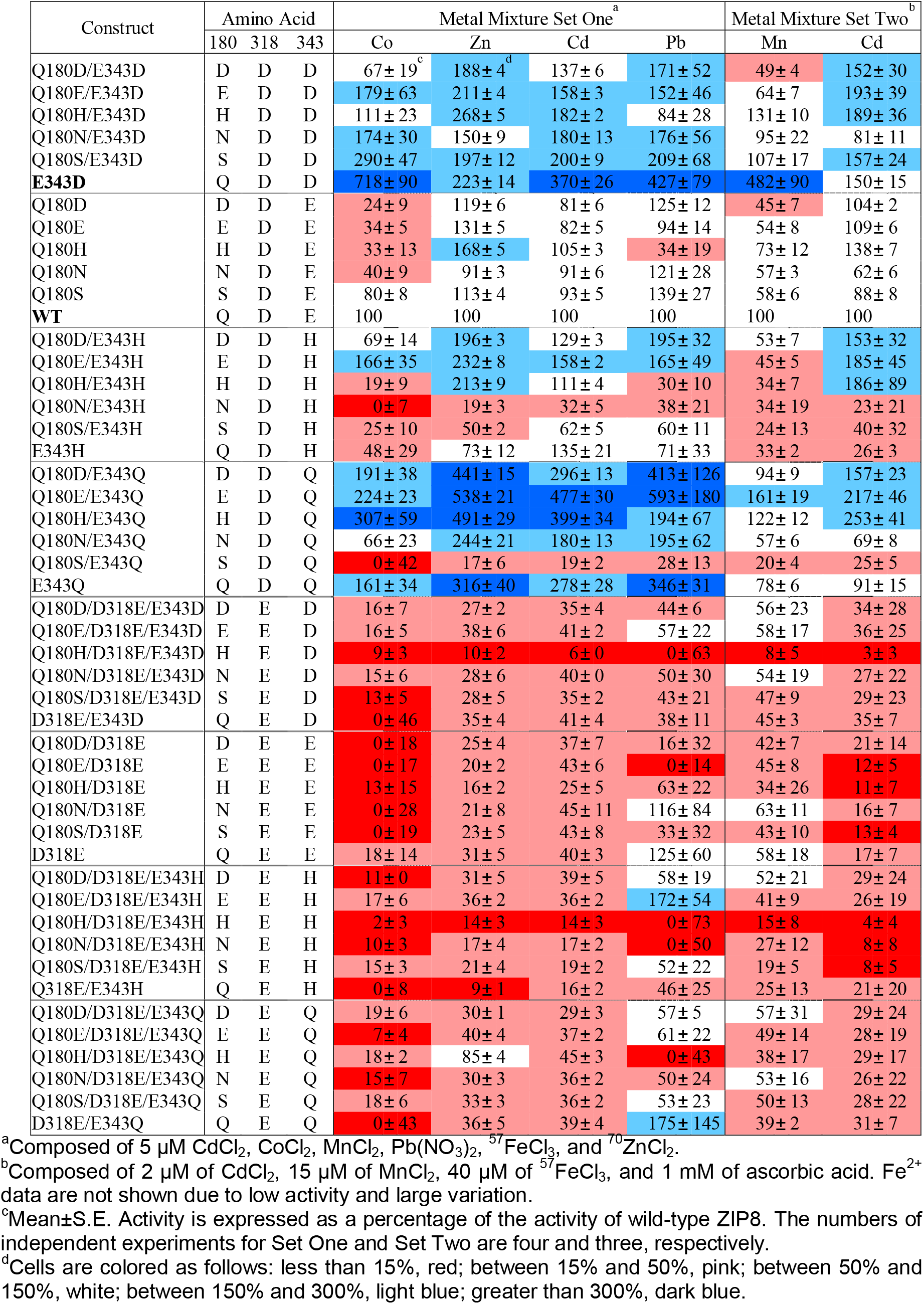
Screening of the ZIP8 construct library against mixtures of metal substrates.

### Metal screen and identification of Pb^2+^ as a substrate of ZIP8

In order to define the border of the substrate spectrum of wild-type ZIP8, we screened a mixture containing a total of eleven metals at the same concentration of 5 µM for each metal, including the known ZIP8 substrates (Zn as ^70^Zn, Fe as ^57^Fe, Mn, Co, and Cd) and those that are not (V as VO^2+^, Ni, Cu, Pb, Au as [AuCl_4_]^-^, and Pt as [PtCl_6_]^2-^), in the presence of 2 mM CaCl_2_ and 1 mM MgCl_2_. Different from our previous studies where a Chelex-treated cell culture medium (DMEM plus 10% fetal bovine serum) was used in the transport assay, we used a chelator-free, serum-free buffer in this work to avoid differential metal binding to serum proteins and small molecule chelators in the cell culture medium. In the transport assay, HEK293T cells transiently expressing ZIP8 (or its variants) were incubated with the buffer containing the metal mixture, and ICP-MS was used to measure the content of each metal in cells, which are expressed as the molar ratio of metal and phosphorus (M/^31^P). Our previous study on ZIP4 showed that using phosphorus to calibrate metal uptake improved data precision by levelling out variations in cell number between samples.^36^ As expected, the uptakes of the known ZIP8 substrates, including Zn^2+^, Cd^2+^, Mn^2+^ and Co^2+^, into the cells expressing ZIP8 were significantly higher than the cells transfected with an empty vector (**Figure 2A**), indicative of the transport activities for these metals. No transport activity of ^57^Fe, a stable Fe isotope with a low natural abundance (2.1%), was detected because ascorbic acid, a reducing agent that would reduce Fe^3+^ to Fe^2+^, is incompatible with several metals (Cu^2+^, Pb^2+^, VO^2+^, [AuCl_4_]^-^, and [PtCl_6_]^2-^) and was therefore excluded from the metal mixture. The lack of ^57^Fe^3+^ transport activity is consistent with the notion that Fe^3+^ is not a ZIP8 substrate. Unexpectedly, the transport activity for Pb^2+^ was detected as the M/^31^P ratio for the cells expressing ZIP8 was significantly higher than that of the cells transfected with an empty vector (**Figure 2A**). In the metal competition assay, 10-fold excess Zn^2+^ almost abolished Pb^2+^ uptake, but 10-fold excess Pb^2+^ did not significantly reduce Zn^2+^ transport (**Figure 2B**), suggesting that the affinity for Pb^2+^ is much lower than that for Zn. It has been shown that only very high concentration of Pb^2+^ at a 30-fold excess was able to significantly block ZIP8-mediated Zn^2+^ and Cd^2+^ transport.^27^ The low affinity for Pb^2+^ was then supported by a kinetic study shown in a later section. As the radius of Pb^2+^ (119 pm) is much larger than Zn^2+^ (74 pm), identification of Pb^2+^ as a substrate indicates that the transport pathway of ZIP8 is flexible enough to allow the transport of metal ions with diverse sizes. Even so, the inability of wild-type ZIP8 to transport VO^2+^, Ni^2+^, Cu^2+^, [AuCl_4_]^-^, and [PtCl_6_]^2-^ indicates that despite its promiscuity, ZIP8 still transports metals in a selective manner.

**Figure 2.**
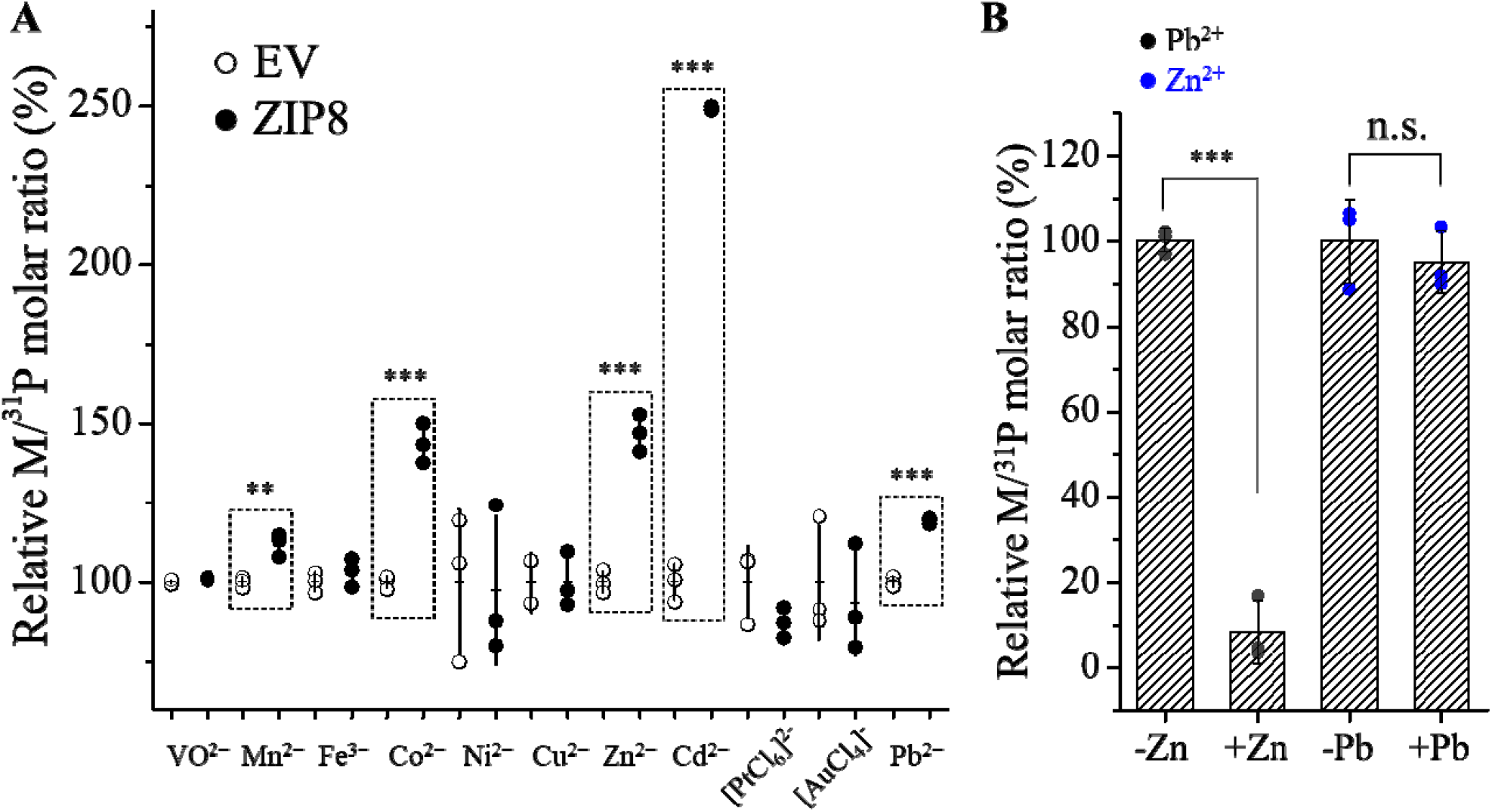
Detection of ZIP8 transport activities for a panel of metal ions. (**A**) Screening for cellular metal content after 30 min of uptake using ICP-MS. Cells were incubated in a solution containing a mixture of eleven metals. The relative M/^31^P molar ratio of the ZIP8 group is expressed as the percentage of the M/^31^P molar ratio of the empty vector group. Each symbol represents the result of one of three replicates for each condition. The horizontal bar is the mean of three replicates and the vertical bar shows 1±S.D. Two-tailed Student’s *t* test was performed to examine statistical significance. ^**^: *P*<0.01; ^***^: *P*<0.001. The *P* values for Mn^2+^, Co^2+^, Zn^2+^, Cd^2+^, and Pb^2+^ are 0.006, 0.0003, 0.0003, 2×10^−6^, and 4×10^−5^, respectively. (**B**) Transport of Pb^2+^ (or Zn^2+^) at 5 µM in the absence and presence of Zn^2+^ (or Pb^2+^) at 50 µM. The shown data are from one of two independent experiments with similar results.

### Library screen to identify variants with altered transport properties

Because ICP-MS can quantify multiple elements in a single sample, it is particularly useful for studying substrate specificity of metal transporters, as the relative transport rate is not affected by the variations in expression of the transporter of interest among different samples. In this work, the 48 constructs in the library were screened against a mixture of metal substrates – Zn^2+^, Cd^2+^, Mn^2+^, Co^2+^, Fe^2+^, and Pb^2+^ (Set One). However, due to the competition between the substrates, the activities of Mn^2+^ and Fe^2+^ couldn’t be consistently measured, suggestive of their relative low transport activities when compared to other metals. To address this problem, a second metal mixture composed of Mn^2+^, Fe^2+^, and Cd^2+^ (Set Two) was set up to screen the library. The molar ratios of Mn^2+^, Fe^2+^, and Cd^2+^ have been optimized so that their activities can be detected for wild-type ZIP8 (**Figure S1**). The M/^31^P ratios of the variants were calculated and expressed as a percentage of the M/^31^P ratio for wild-type ZIP8, and the results from 3-4 independent experiments are summarized in **Table 1**. The expression of ZIP8 and its variants was detected by Western blot (**Figure S2**). The Fe^2+^ uptake data are not listed in **Table 1** because the readings for many variants were less reproducible with larger deviations than those for other metals, likely due to the weak Fe^2+^ transport activity even under the optimized conditions. Kinetic studies of Fe^2+^ transport of wild-type ZIP8 and a selected variant (E343D) are described in the next section. Analysis of the data in **Table 1** revealed the following findings:

i. The D318E mutation is detrimental to the transport of nearly all metals for all variants. For most D318E-containing variants, the abolished activity was not due to altered expression according to the results of the Western blot experiments (**Figure S2**).
ii. The mutations on E343 have different impacts on transport activity. The E343D mutation alone or in combination with Q180 mutations led to increased activities for most metals. Similarly, the E343Q mutation alone or in combination with Q180 mutations (except for Q180S) favored the transport of most metals. In contrast, the variants containing the E343H mutation exhibited very different activities. Among them, the double variant Q180H/E343H showed a selectively increased activity for Zn^2+^, consistent with our previous study.^31^ Q180H alone also improved selectivity for Zn^2+^, and the effect was strengthened when combined with E343H, as reported previously.^31^
iii. There are positive correlations between the activities of any two metals (**Figure 3**), when the E343D variant and the variants containing the D318E mutation are excluded. The greatest correlation was found to be between Zn^2+^ and Cd^2+^, which is consistent with their similar chemical properties and therefore it would be most challenging to create variants to selectively transport one while exclude the other. In contrast, the poorest correlations between Mn^2+^ and Cd^2+^ and between Co^2+^ and Pb^2+^ imply better chances to separate the activities in each metal pair. While the slopes of the Zn-involved linear regression were all smaller than one (**Figure 3**, first row), indicating that most of the mutations at the selectivity filter is to increase the preference for Zn^2+^ over other metals, the E343D variant is an apparent outlier because it shows a preference for non-zinc metals.

**Figure 3.**
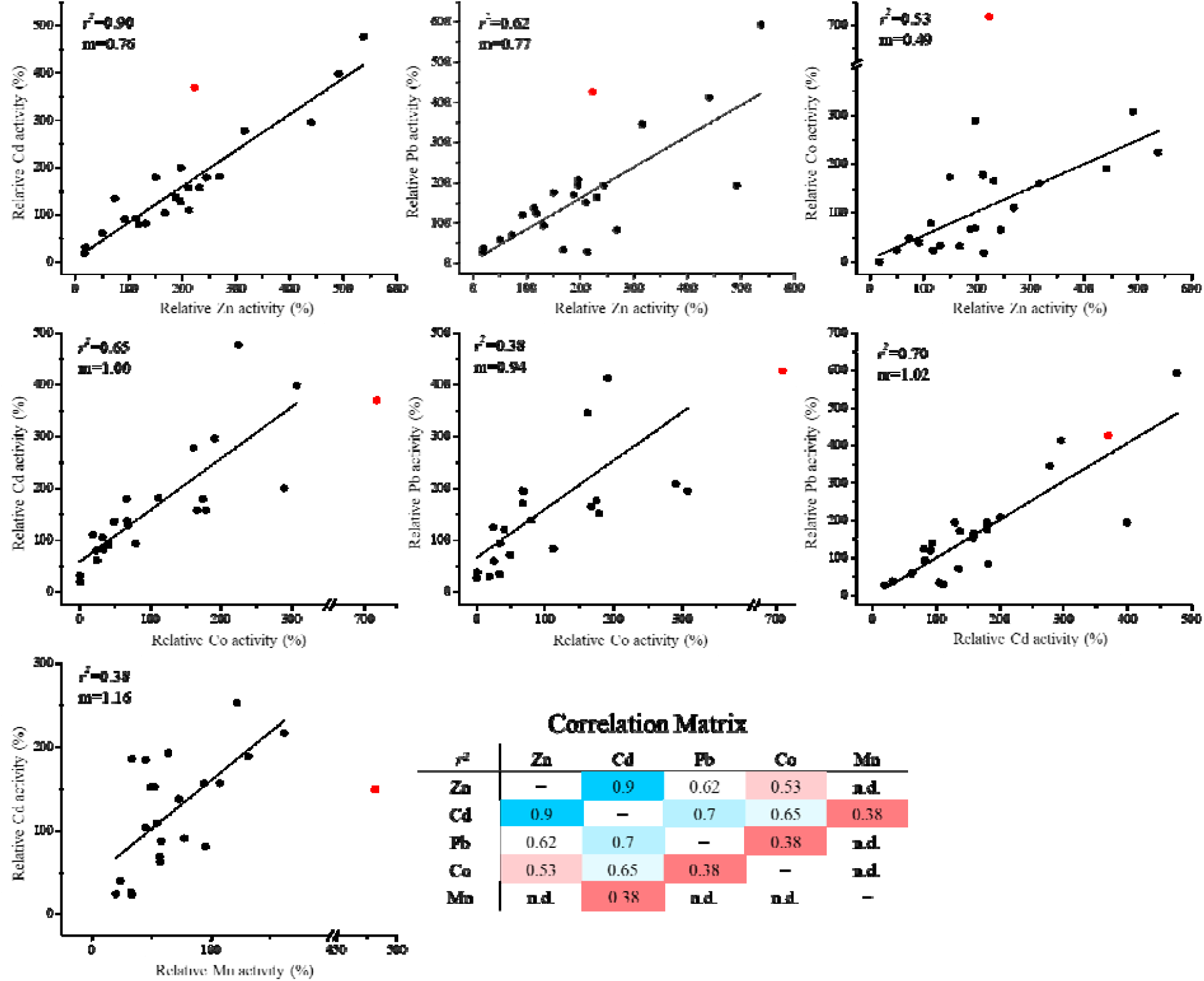
Correlation analysis of the activities of different metal substrates of 24 constructs of ZIP8. Activities are expressed as percentages of the corresponding activities of wild-type ZIP8. Only the variants without the D318E substitution were analyzed due to the strong detrimental effects of the D318E mutation on the activities for nearly all metals tested. The E343D variant (red solid circle) was excluded from linear regression because it appears to be an outlier in several cases. The activities for Zn^2+^, Cd^2+^, Pb^2+^, and Co^2+^ obtained in metal mixture Set One and the activities for Mn^2+^ and Cd^2+^ obtained in metal mixture Set Two were applied to correlation analysis. The *r*^2^ values are summarized in the correlation matrix.

### Kinetic study of the E343D variant

As the E343D variant exhibited transport properties different from the other variants, we performed kinetic studies to understand the mechanism of its broadly increased activities and altered substrate preference. Kinetic studies were performed on single metals to avoid interference between competing substrates. The molar ratios of M/^31^P obtained from ICP-MS were plotted against the indicated metal concentrations, following by curve fitting using the Michaelis-Menten model. As shown in **Figures 4A & 4B**, the results of the kinetic study consistently showed that the E343D variant exhibited a significant increase in *V*_max_ for Zn^2+^, Cd^2+^, Mn^2+^, and Fe^2+^ by 2-3 folds, while the *K*_M_ values for these metals were also increased. Although the E343D mutation did not significantly change the order of the *V*_max_/*K*_M_ values, i.e. Zn^2+^∼Cd^2+^∼Co^2+^>Mn^2+^>Fe^2+^, the variant showed enhanced transport of Cd^2+^ and Fe^2+^ more than the other metals, indicative of an altered substrate preference. We noticed that the largely preferred transport for Co^2+^ and Mn^2+^ of this variant, as indicated in the screening of metal mixtures (**Table 1 & Figure 3**), cannot be explained by the measured kinetic parameters. This discrepancy suggests that the metal substrates do not simply compete for the single high-affinity transport site, and therefore the assumption that the relative transport rate obtained from the experiments using a substrate mixture accurately reflects the ratios of the *V*_max_/*K*_M_ values determined using individual substrates does not hold any more. Indeed, the involvement of the selectivity filter, a weak but critical metal binding site, may complicate the kinetic analysis. Allostery between Zn^2+^ and Mn^2+^ was also noticed in our previous study on a ZIP8 variant.^31^ For Pb^2+^, we were not able to obtain the saturable kinetic parameters: the dose-response curves were almost linear, suggestive of poor binding and consistent with the results of the competition experiment (**Figure 2B**). Even so, it is clear that the E343D variant transports Pb^2+^ faster than the wild-type transporter. The increased transport activity was not due to a higher expression level of the variant. Western blot and flow cytometry data showed that the E343D variant actually has a lower total expression level but a similar cell surface expression level when compared to wild-type ZIP8 (**Figures 4C & S4**). Therefore, the *k*_cat_ value of this variant is approximately 2-3 times higher than the wild-type transporter, as indicated by the relative *V*_max_ values (**Figure 4B**). Taken together, the kinetic studies showed that the E343D variant transport all the metals tested more efficiently with Cd^2+^ and Fe^2+^ being the most enhanced, while the apparent affinities for metals were all reduced.

**Figure 4.**
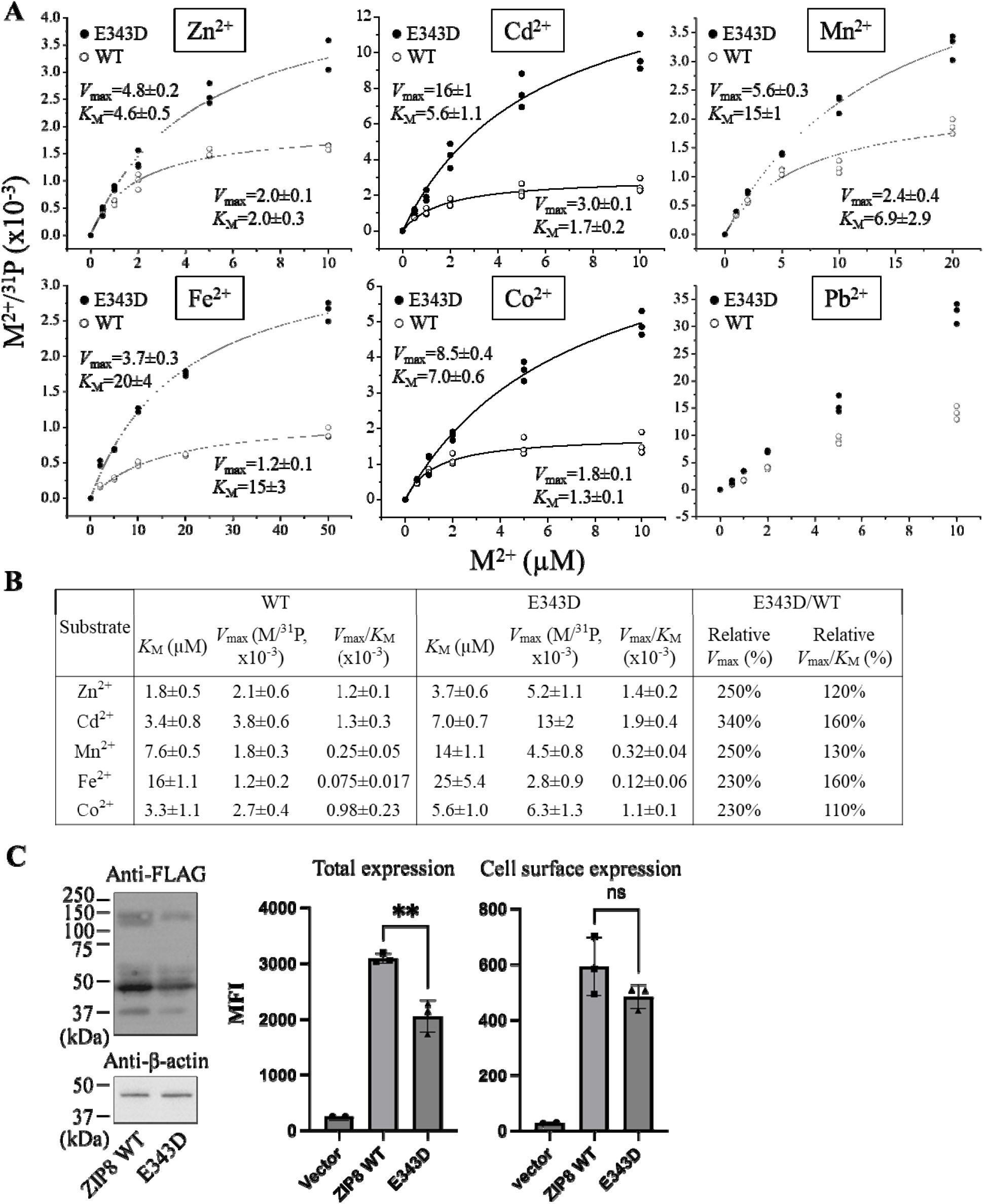
Kinetic studies of the E343D variant in comparison with wild-type ZIP8. (**A**) Transport assay of six metal substrates. The molar ratios of the metals and ^31^P were plotted against the concentrations of the metals. The shown data are from one of the three independent experiments with similar results. Curve fittings were conducted using the Michaelis-Menten model except for the curves for Co^2+^ and Pb^2+^. Each data point represents the result of one of the three biological replicates for one condition. The uncertainties shown in the figure are the standard errors of curve fitting. (**B**) Summary of the kinetic parameters from the three independent experiments. *V*_max_ and *V*_max_/*K*_M_ are expressed as the percentage relative to wild-type ZIP8 measured in the same experiment. The data are expressed as mean±S.E. (n=3). N.A.: not available. (**C**) Comparison of the expression levels of wild-type ZIP8 and the E343D variant. *Left*: total expression of the FLAG-tagged ZIP8 and E343D variant detected by Western blot. The total expression level of the variant is about half of that of wild-type ZIP8. *Right*: total expression and cell surface expression measured by flow cytometry using an anti-FLAG antibody and an Alexa Fluor 568-labeled anti-mouse IgG. MFI: mean fluorescence intensity. Cells were fixed with and without permeabilization prior to staining for detection of the total and cell surface expression, respectively. Two-tailed Student’s *t* test was used to examine statistical significance. **: *P*<0.01. Data processing is illustrated in Figure S4.

### New transport activities for non-substrate metals

Since the E343D variant exhibited a significantly increased transport efficiency and an altered substrate preference, we wondered if this variant could transport metals that are not the substrates for wild-type ZIP8. We then tested two divalent ions, VO^2+^ which is a diatomic ion with a dimension greater than any known ZIP8 substrates, and Cu^2+^, which has been reported to be transported by various ZIPs^41, 42^ but not by ZIP8 in this study (**Figure 2**). Of great interest, the E343D variant exhibited transport activities for both metals (**Figure 5**), even though the activities are marginal and the dose-response curves are not saturable, suggestive of weak binding. These results indicate that the E343D variant has an expanded substrate spectrum and thus becomes more promiscuous than wild-type ZIP8.

**Figure 5.**
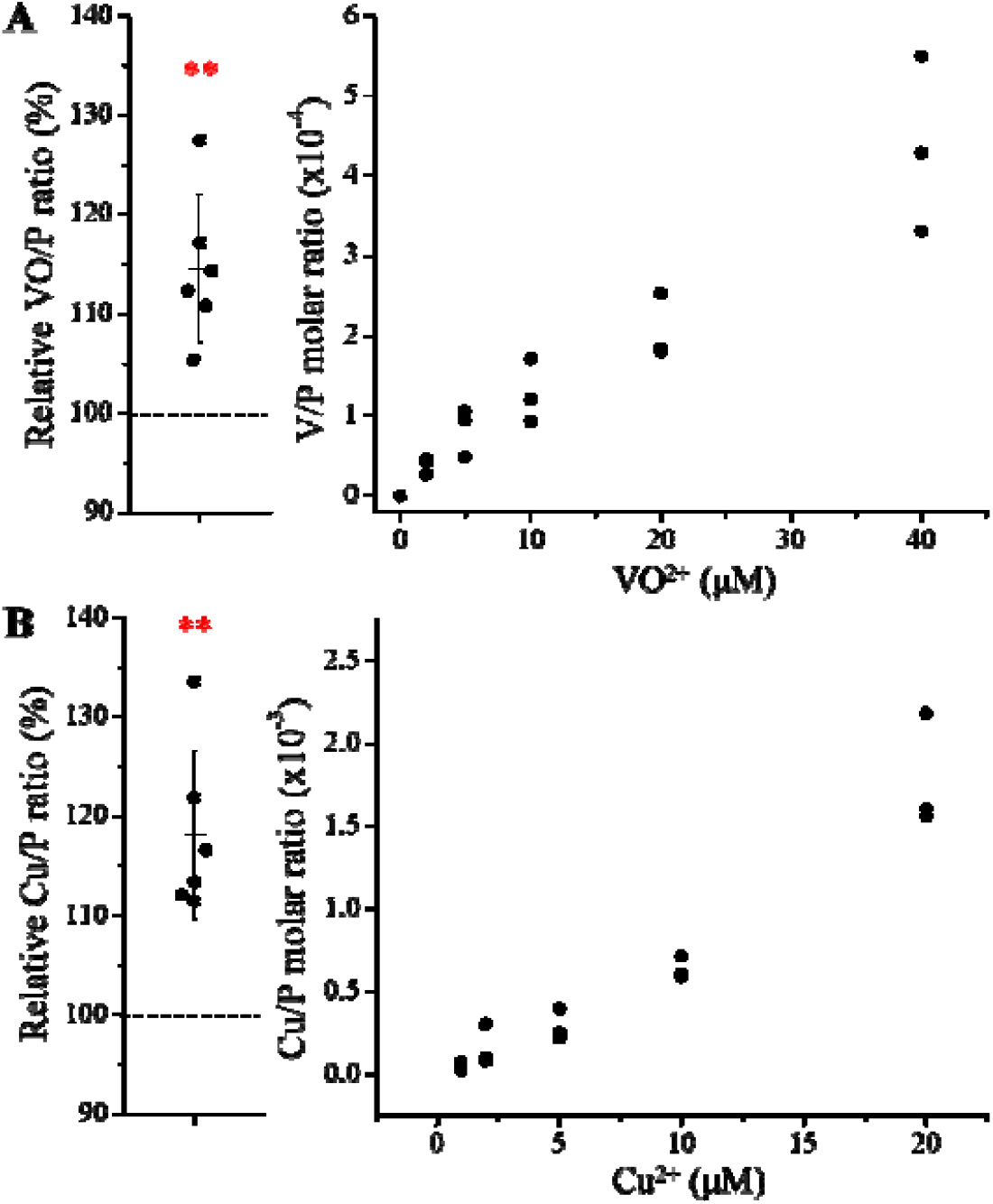
New transport activities of the E343D variant for VO^2+^ (**A**) and Cu^2+^ (**B**). The results of six independent experiments with cells incubated with 5 μM M^2+^ are shown on the left. The dashed lines indicate the normalized M^2+^ uptake by the cells transfected with an empty vector (set as 100%). Statistical analyses were performed using two-tailed Student’s *t* test. The *P* values for VO^2+^ and Cu^2+^ are 0.0048 and 0.0033, respectively. **: *P*<0.01. The dose-response curves for VO^2+^ and Cu^2+^ from one of two independent experiments with similar results are shown on the right with three replicates tested for each condition.

## Discussion

Engineering of metal transporters for enrichment or exclusion of specific metals from living organisms has great potential for a variety of applications. For example, expressing importers specific for beneficial metals in crop roots may help to address mineral deficiency problems in populations,^10, 11^ while carefully tuning the substrate spectra of these metal importers may provide a solution to avoid heavy metal accumulation in edible parts of crops by eliminating activities for toxic metals while reserving those for beneficial ones.^16, 17^ In engineered plants/algae, overexpression of toxic metal-specific importers can remove heavy metals from the environment,^12, 13^ and this strategy can also be used to harvest low abundance precious metals from souroundings.^14, 15^ However, these prospects still face several significant obstacles. First, the *d*-block metals that are the primary targets in these applications have similar chemical properties, making it difficult to create metal transporters with high selectivity. Second, our understanding of the transport mechanism and substrate specificity of *d*-block metal transporters is still in its infancy, and the poor knowledge of the dynamic *d*-block metal-protein interactions during metal translocation hinders rational engineering. Third, the lack of high-throughput assays for most metal transporters impedes the application of directed evolution to metal transporter engineering. In this work, we focused on the selectivity filter at the pore entrance of ZIP8,^31^ a promiscuous *d*-block divalent transporter, to explore the potential of changing the transport properties, especially the substrate preference, for subsequent large-scale metal transporter engineering. Our results indicated that both the transport activity and substrate specificity of ZIP8 can be altered over a wide range (**Table 1, Figures 4&5**), demonstrating the feasibility of tuning the function of a metal transporter by targeting the key residues selected based on prior knowledge of structure and mechanism.

By altering the three residues that form the selectivity filter, we generated a library consisting of 48 constructs and screened them using ICP-MS to measure the transport activities for multiple metals simultaneously. Among the variants, the E343D variant is particularly interesting because of the significantly enhanced transport activity and altered substrate preference. Notably, the position of E343 is not occupied by aspartate in any of the known ZIP,^35^ even though glutamate and aspartate have very similar properties. The kinetic study showed that the E343D mutation increased the transport rate (*V*_max_) at the cost of affinity (**Figure 4**), which is similar to the affinity-activity trade-off for enzymes.^43^ Since the increase in *V*_max_ is greater than that in the apparent *K*_M_, the E343D variant exhibited enhanced transport efficiencies (*V*_max_/*K*_M_) for all tested metals with the transport of Cd^2+^ and Fe^2+^ increasing most. Structurally, as E343 participates in both the selectivity filter and the transport site, the increased transport efficiency caused by the E343D substitution suggests that a shorter side chain may allow metals to more rapidly reach the transport site due to a larger pore size of the selectivity filter and/or an expedited release from the transport site due to a longer distance between the bound metal and the carboxylic acid group and thus a weaker interaction. These effects on the structures of the selectivity filter and the transport site may also explain why the E343D variant becomes more promiscuous than wild-type ZIP8 (**Figure 5**). The increased promiscuity makes this variant a better choice than the wild-type ZIP8 for later directed evolution study,^44, 45^ and this variant may also be used in the scenarios where a ZIP8 with an enhanced transport activity is preferred.

In contrast to the activity-enhancing E343D mutation, the D318E substitution drastically reduced the transporter activity across the entire spectrum of metal substrates. This may be attributed to a smaller pore size of the selectivity filter and a reduced volume of the transport site. However, the activity reduction associated with the D318E mutation could not be rescued by the E343D substitution, despite the fact that these two residues in the WT structure face each other in the selectivity filter and the transport site (**Figure 1**). The lack of epistatic effects of contacting residues suggests that D318 and E343 have non-overlapping functions. In fact, the variants with the same amino acid composition at the selectivity filter but different distribution among the three key residues often did not show the same activity and substrate preference (**Figure S3**). We conclude that it is not simply the amino acid composition but also the side chain positions in the selectivity filter and the transport site that are important for transport activity and substrate preference.

Importantly, our results also reveal a remarkable potential of the ZIP fold to bind and transport metal ions with diverse physicochemical properties. It is unexpected that Pb^2+^, a soft, non-*d*-block metal ion is able to translocate through ZIP8. This is remarkable because Pb^2+^ has an ionic radius 23% larger than Cd^2+^ (97 pm), the previously largest ZIP substrate. (**Figure 2**). A recent study reported Cu^2+^ transport activities for several human ZIPs expressed in HEK293 cells, including ZIP8.^42^ In fact, we detected a marginal Cu^2+^ transport activity for the E343D variant (**Figure 5**) but not for wild-type ZIP8 (**Figure 2**). We noticed that there are differences in experimental settings, such as the absence of EDTA or other metal chelators in wash buffer used in that work and these may account for some differences. We also find that VO^2+^ is not a substrate for wild-type ZIP8, but the new activity towards this diatomic ion was detected for the E343D variant (**Figure 5**). The new transport activities for Pb^2+^, VO^2+^ and Cu^2+^, together with the highly variable activity and substrate preference observed in the variant library, demonstrate that the ZIP fold, which is uniquely present and conserved in the ZIP family, has a remarkable plasticity and amenability, enabling it to transport a wide range of metal ions. This flexibility suggests that it could serve as a versatile scaffold for creating novel metal transporters with significantly expanded or tailored substrate specificity. For instance, the ZIP8 variants with increased substrate specificity toward one metal over others may be valuable tools to dissect the multi-functions of ZIP8 in physio-pathological scenarios involving different metals.^46-51^

## Materials and Methods

### Gene, plasmids, and reagents

The complementary DNA of human ZIP8 (GenBank access number: BC012125) from Mammalian Gene Collection were purchased from GE Healthcare. The ZIP8 construct consists of the N-terminal signal peptide of ZIP4 (amino acid residues 1-22) followed by a GSGS linker and a FLAG tag, and the ZIP8 coding sequence (residue 23-460). Site-directed mutagenesis of ZIP8 was conducted using QuikChange mutagenesis kit (Agilent, Cat#600250). All mutations were verified by DNA sequencing. ^70^ZnO was purchased from American Elements (ZN-OX-01-ISO.070I, Lot#1871511028-401) and ^57^FeCl_3_ was purchased from Sigma-Aldrich (790427, Lot#MBBD4771). 30 mg of zinc oxide powder was dissolved in 5 ml of 1 M HCl and then diluted with ddH_2_O to make the stock solution at the concentration of 50 mM. ^57^FeCl_3_ was dissolved in 1 M HCl to final concentration of 100 mM. The ^70^Zn sample was certified as 72% abundance and ^57^Fe was 99.9%. Other reagents were purchased from Sigma-Aldrich or Fisher Scientific.

### Cell culture, transfection, and Western blot

Human embryonic kidney cells (HEK293T, ATCC, Cat#CRL-3216) were cultured in Dulbecco’s modified eagle medium (DMEM, Thermo Fisher Scientific, Invitrogen, Cat#11965092) supplemented with 10% (v/v) fetal bovine serum (FBS, Thermo Fisher Scientific, Invitrogen, Cat#10082147) and 1% Antibiotic-Antimycotic solution (Thermo Fisher Scientific, Invitrogen, Cat# 15240062) at 5% CO_2_ and 37°C. Cells were seeded on the polystyrene 24-well trays (Alkali Scientific, Cat#TPN1024) coated with poly-D-lysine (Corning, Cat#354210) for 16 h and transfected with 0.8 μg DNA/well using lipofectamine 2000 (Thermo Fisher Scientific, Invitrogen, Cat# 11668019) in DMEM with 10% FBS.

For Western blot, samples were mixed with the SDS sample loading buffer and heated at 37°C for 20 min before loading on SDS-PAGE gel. The proteins separated by SDS-PAGE were transferred to PVDF membranes (Millipore, Cat#PVH00010). After blocking with 5% (w/v) nonfat dry milk, the membranes were incubated with anti-FLAG antibody (Agilent, Cat# 200474-21) at 1:3000 or anti β-actin (Cell Signaling, Cat# 4970S) at 1:2500 at 4°C overnight, which were detected with HRP-conjugated goat anti-rat immunoglobulin-G at 1:5000 dilution (Cell Signaling Technology, Cat# 7077S) or goat anti-rabbit immunoglobulin-G at 1:3000 dilution (Cell Signaling Technology, Cat# 7074S) respectively using the chemiluminescence reagent (VWR, Cat#RPN2232). The images of the blots were taken using a Bio-Rad ChemiDoc Imaging System.

### Metal transport assay

Twenty hours post transfection, cells were washed with a chelator-free, serum-free incubation buffer (10 mM Hepes, 142 mM NaCl, 5 mM KCl, 10 mM glucose, pH 7.3) followed by incubation with the incubation buffer plus metals for 30 min. To screen substrate and non-substrate metals for wild-type ZIP8 (**Figure 1**), the metal mixture contained the following compounds: 2 mM CaCl_2_, 1 mM MgCl_2_, 5 µM for CdCl_2_, CoCl_2_, MnCl_2_, Pb(NO_3_)_2_, ^57^FeCl_3_, ^70^ZnCl_2_, VoCl_2_, NiCl_2_, CuCl_2_, AuCl_3_ (as [AuCl_4_]^-^), and PtCl_4_ (as [PtCl_6_]^2-^). Given the *K*_sp_ of PbCl_2_ (1.6×10^−5^ at 20°C), Pb^2+^ at 5-50 µM in the above buffer is soluble. To screen variants against metal substrates, metal mixture Set One is composed of 2 mM CaCl_2_, 1 mM MgCl_2_, 5 µM CdCl_2_, CoCl_2_, MnCl_2_, Pb(NO_3_)_2_, ^57^FeCl_3_, and ^70^ZnCl_2_ in the incubation buffer. The metal mixture Set Two is composed of 2 mM CaCl_2_, 1 mM MgCl_2_, 2 µM of CdCl_2_, 15 µM of MnCl_2_, 40 µM of ^57^FeCl_3_ and 1 mM of ascorbic acid in the incubation buffer. After incubation at 37°C for 30 min, the 24-well plates were put on ice and an equal volume of the ice-cold washing buffer containing 1 mM EDTA was added to the cells to terminate metal uptake, followed by three times of washing with ice-cold wash buffer before cell lysis.

For kinetic studies, cells were grown and transfected in the same way as in the metal transport assay. Briefly, twenty hours post transfection, cells were washed with the same incubation buffer, followed by incubation with the buffer plus metals for 30 min. All metal solutions contained 2 mM CaCl_2_ and 1 mM MgCl_2_, with different concentrations of metals as indicated. 1 mM ascorbic acid was included when the transport assay for Fe^2+^ was conducted. After incubation at 37°C for 30 minutes, 24-well plates were put on ice and an equal volume of the ice-cold washing buffer containing 1 mM EDTA was added to cells to terminate metal uptake, followed by three times of washing with ice-cold wash buffer before cell lysis in nitric acid.

### ICP-MS experiment

The ICP-MS experiments were conducted as described.^31^ All standards, blanks, and cell samples were prepared using trace metal grade nitric acid (70%, Fisher chemical, Cat# A509P212), ultrapure water (18.2 MΩ·cm @ 25 °C), and metal free polypropylene conical tubes (15 and 50 mL, Labcon, Petaluma, CA, USA). For cell samples in polystyrene 24-well cell culture plates, 200 µl of 70% trace nitric acid was added to allow for initial sample digestion. Following digestion, 150 µl of the digested product was transferred into metal free 15 mL conical tubes. For liquid samples, 50 µl of liquid samples were added to metal free conical tubes followed by addition of 150 µl of 70% trace nitric acid. All cell and liquid samples were then incubated at 65°C in a water bath for one hour followed by dilution to 5 ml using ultrapure water. These completed ICP-MS samples were then analyzed using the Agilent 8900 Triple Quadrupole ICP-MS (Agilent, Santa Clara, CA, USA) equipped with the Agilent SPS 4 Autosampler, integrate sample introduction system (ISiS), x-lens, and micromist nebulizer. Daily tuning of the instrument was accomplished using manufacturer supplied tuning solution containing Li, Co, Y, Ce, and Tl. Global tune optimization was based on optimizing intensities for ^7^Li, ^89^Y, and ^205^Tl while minimizing oxides (^140^Ce^16^O/^140^Ce < 1.5%) and doubly charged species (^140^Ce++/^140^Ce+ < 2%). Following global instrument tuning, gas mode tuning was accomplished using the same manufacturer supplied tuning solution in KED mode (using 100% UHP He, Airgas). Specifically, intensities for ^59^Co, ^89^Y, and ^205^Tl were maximized while minimizing oxides (^140^Ce^16^O/^140^Ce < 0.5%) and doubly charged species (^140^Ce++/^140^Ce+ < 1.5%) with short term RSDs < 3.5%. ICP-MS standards were prepared from a stock solution of NWU-16 multi-element standard (Inorganic Ventures, Christiansburg, VA, USA) that contains the metals studied in this work that were diluted with 3% (v/v) trace nitric acid in ultrapure water to a final element concentration of 1000, 500, 250, 125, 62.5, 31.25, and 0 (blank) ng/g standard. Internal standardization was accomplished inline using the ISIS valve and a 200 ng/g internal standard solution in 3% (v/v) trace nitric acid in ultrapure water consisting of Bi, In, ^6^Li, Sc, Tb, and Y (IV-ICPMS-71D, Inorganic Ventures, Christiansburg, VA, USA). ^6^Li, ^45^Sc, and ^89^Y were used for internal standardization. Continuing calibration blanks (CCBs) were run every 10 samples and a continuing calibration verification standard was analyzed at the end of every run for a 90-110% recovery.

### Flow cytometry experiment

To detect the total expression level, the cells were fixed with 4% formaldehyde at room temperature for 15 min, blocked and permeabilized for 1 h with Dulbecco’s phosphate-buffered saline (DPBS) containing 2% BSA and 0.1% Triton X-100, and then incubated with anti-FLAG antibody (Sigma, F3165) at 1:500 in 2% BSA in PBST containing 0.1% Tween-20 (ThermoFisher Scientific, Cat# 85113) for 2 hours on ice. After the unbound antibodies were removed by wash using DPBS, the cells were incubated with an AlexaFluor 568-conjugated goat anti-mouse antibody (Invitrogen, CAT#A-11004) at 1: 1000 diluted in DPBS with 2% BSA for 1 h. After three time wash with a FACS buffer (2% BSA, 1 mM EDTA, 20 mM HEPES in DPBS without Ca^2+^ and Mg^2+^), the cells were resuspended by FACS buffer for flow cytometry analysis. To detect the surface expression level, the procedure was the same except that Triton X-100 and Tween-20 were excluded. The cells were applied to a ThermoFisher Attune Cytpix flow cytometer. 10,000 cells were assessed for each condition. The single cells were selected (**Figure S4**) and then analysed in the YL-1 channel to detect the fluorescence of AlexaFluor-568. The mean fluorescence intensity (MFI) of each population was calculated and the histogram charts were generated to compare the signals from each group (**Figure S4**). All the data processing was performed in FlowJo 10.8.1 and GraphPad Prism 10.

## Statistics

We assumed a normal distribution of the samples and significant difference were examined using two-tail Student’s *t* test. Uncertainties shown in figures are reported as standard deviation (S.D.) or standard error (S.E.).

## Data availability

All raw and processed data reported in the main text and SI are available upon request. The structure shown in Figures 5, 6, and S1 is retrieved from PDB with the accession code 5TSB.

## Acknowledgments

This work is supported by National Institutes of Health R35GM140931 (to J. H.) and P41GM135018, R01GM115848, and R01GM038784 (to T.V.O.). The data presented herein were obtained using instrumentation in the MSU Flow Cytometry Core Facility. The facility is funded in part through the financial support of Michigan State University’s Office of Research & Innovation and Colleges of Osteopathic Medicine, Human Medicine, Veterinary Medicine, Natural Sciences, and Engineering. The Attune CytPix is supported by the Equipment Grants Program, award #2022-70410-38419, from the U.S. Department of Agriculture, National Institute of Food and Agriculture. The content is solely the responsibility of the authors and does not necessarily represent the official views of the National Institutes of Health.The content is solely the responsibility of the authors and does not necessarily represent the official views of the National Institutes of Health.

## Author contributions

J. H., Y.J., M.N., T.W., K.M., and T.V.O. conceived the project and designed the experiments, Y.J., M.N., T.W., and K.M. conducted the experiments; Y.J., M.N., T.W., K.M., T.V.O., and J. H. analyzed the data and wrote the manuscript.

## References

1. Maret, W., Zinc and human disease. Met Ions Life Sci 2013, 13, 389–414.

2. Erikson, K. M.; Aschner, M., Manganese: Its Role in Disease and Health. Met Ions Life Sci 2019, 19.

3. Abbaspour, N.; Hurrell, R.; Kelishadi, R., Review on iron and its importance for human health. J Res Med Sci 2014, 19 (2), 164–74.

4. Yamada, K., Cobalt: its role in health and disease. Met Ions Life Sci 2013, 13, 295–320.

5. Zamble, D., Introduction to the Biological Chemistry of Nickel. Rsc Metallobio Ser 2017, 10, 1–11.

6. Festa, R. A.; Thiele, D. J., Copper: an essential metal in biology. Curr Biol 2011, 21 (21), R877–83.

7. Mendel, R. R.; Bittner, F., Cell biology of molybdenum. Biochim Biophys Acta 2006, 1763 (7), 621–35.

8. Nelson, N., Metal ion transporters and homeostasis. EMBO J 1999, 18 (16), 4361–71.

9. Ma, Z.; Jacobsen, F. E.; Giedroc, D. P., Coordination chemistry of bacterial metal transport and sensing. Chem Rev 2009, 109 (10), 4644–81.

10. Narayanan, N.; Beyene, G.; Chauhan, R. D.; Gaitan-Solis, E.; Gehan, J.; Butts, P.; Siritunga, D.; Okwuonu, I.; Woll, A.; Jimenez-Aguilar, D. M.; Boy, E.; Grusak, M. A.; Anderson, P.; Taylor, N. J., Biofortification of field-grown cassava by engineering expression of an iron transporter and ferritin. Nat Biotechnol 2019, 37 (2), 144–151.

11. Stanton, C.; Sanders, D.; Kramer, U.; Podar, D., Zinc in plants: Integrating homeostasis and biofortification. Mol Plant 2022, 15 (1), 65–85.

12. Yang, Z.; Yang, F.; Liu, J. L.; Wu, H. T.; Yang, H.; Shi, Y.; Liu, J.; Zhang, Y. F.; Luo, Y. R.; Chen, K. M., Heavy metal transporters: Functional mechanisms, regulation, and application in phytoremediation. Sci Total Environ 2022, 809, 151099.

13. Kozminska, A.; Wiszniewska, A.; Hanus-Fajerska, E.; Muszynska, E., Recent strategies of increasing metal tolerance and phytoremediation potential using genetic transformation of plants. Plant Biotechnol Rep 2018, 12 (1), 1–14.

14. Navarro, C. A.; von Bernath, D.; Jerez, C. A., Heavy metal resistance strategies of acidophilic bacteria and their acquisition: importance for biomining and bioremediation. Biol Res 2013, 46 (4), 363–71.

15. Antonucci, I.; Gallo, G.; Limauro, D.; Contursi, P.; Ribeiro, A. L.; Blesa, A.; Berenguer, J.; Bartolucci, S.; Fiorentino, G., Characterization of a promiscuous cadmium and arsenic resistance mechanism in Thermus thermophilus HB27 and potential application of a novel bioreporter system. Microb Cell Fact 2018, 17 (1), 78.

16. He, G.; Tian, W.; Qin, L.; Meng, L.; Wu, D.; Huang, Y.; Li, D.; Zhao, D.; He, T., Identification of novel heavy metal detoxification proteins in Solanum tuberosum: Insights to improve food security protection from metal ion stress. Sci Total Environ 2021, 779, 146197.

17. Lee, J.; Bae, H.; Jeong, J.; Lee, J. Y.; Yang, Y. Y.; Hwang, I.; Martinoia, E.; Lee, Y., Functional expression of a bacterial heavy metal transporter in Arabidopsis enhances resistance to and decreases uptake of heavy metals. Plant Physiol 2003, 133 (2), 589–96.

18. Guerinot, M. L., The ZIP family of metal transporters. Biochim Biophys Acta 2000, 1465 (1-2), 190–8.

19. Hu, J., Toward unzipping the ZIP metal transporters: structure, evolution, and implications on drug discovery against cancer. FEBS J 2021, 288 (20), 5805–5825.

20. Jeong, J.; Eide, D. J., The SLC39 family of zinc transporters. Mol Aspects Med 2013, 34 (2-3), 612–9.

21. Eide, D. J., Transcription factors and transporters in zinc homeostasis: lessons learned from fungi. Crit Rev Biochem Mol Biol 2020, 55 (1), 88–110.

22. Kambe, T.; Taylor, K. M.; Fu, D., Zinc transporters and their functional integration in mammalian cells. J Biol Chem 2021, 296, 100320.

23. Rogers, E. E.; Eide, D. J.; Guerinot, M. L., Altered selectivity in an Arabidopsis metal transporter. Proc Natl Acad Sci U S A 2000, 97 (22), 12356–60.

24. He, L.; Girijashanker, K.; Dalton, T. P.; Reed, J.; Li, H.; Soleimani, M.; Nebert, D. W., ZIP8, member of the solute-carrier-39 (SLC39) metal-transporter family: characterization of transporter properties. Mol Pharmacol 2006, 70 (1), 171–80.

25. Liu, Z.; Li, H.; Soleimani, M.; Girijashanker, K.; Reed, J. M.; He, L.; Dalton, T. P.; Nebert, D. W., Cd2+ versus Zn2+ uptake by the ZIP8 HCO3--dependent symporter: kinetics, electrogenicity and trafficking. Biochem Biophys Res Commun 2008, 365 (4), 814–20.

26. Jenkitkasemwong, S.; Wang, C. Y.; Mackenzie, B.; Knutson, M. D., Physiologic implications of metal-ion transport by ZIP14 and ZIP8. Biometals 2012, 25 (4), 643–55.

27. Nebert, D. W.; Galvez-Peralta, M.; Hay, E. B.; Li, H.; Johansson, E.; Yin, C.; Wang, B.; He, L.; Soleimani, M., ZIP14 and ZIP8 zinc/bicarbonate symporters in Xenopus oocytes: characterization of metal uptake and inhibition. Metallomics 2012, 4 (11), 1218–25.

28. Dalton, T. P.; He, L.; Wang, B.; Miller, M. L.; Jin, L.; Stringer, K. F.; Chang, X.; Baxter, C. S.; Nebert, D. W., Identification of mouse SLC39A8 as the transporter responsible for cadmium-induced toxicity in the testis. Proc Natl Acad Sci U S A 2005, 102 (9), 3401–6.

29. Boycott, K. M.; Beaulieu, C. L.; Kernohan, K. D.; Gebril, O. H.; Mhanni, A.; Chudley, A. E.; Redl, D.; Qin, W.; Hampson, S.; Kury, S.; Tetreault, M.; Puffenberger, E. G.; Scott, J. N.; Bezieau, S.; Reis, A.; Uebe, S.; Schumacher, J.; Hegele, R. A.; McLeod, D. R.; Galvez-Peralta, M.; Majewski, J.; Ramaekers, V. T.; Care4Rare Canada, C.; Nebert, D. W.; Innes, A. M.; Parboosingh, J. S.; Abou Jamra, R., Autosomal-Recessive Intellectual Disability with Cerebellar Atrophy Syndrome Caused by Mutation of the Manganese and Zinc Transporter Gene SLC39A8. Am J Hum Genet 2015, 97 (6), 886–93.

30. Park, J. H.; Hogrebe, M.; Gruneberg, M.; DuChesne, I.; von der Heiden, A. L.; Reunert, J.; Schlingmann, K. P.; Boycott, K. M.; Beaulieu, C. L.; Mhanni, A. A.; Innes, A. M.; Hortnagel, K.; Biskup, S.; Gleixner, E. M.; Kurlemann, G.; Fiedler, B.; Omran, H.; Rutsch, F.; Wada, Y.; Tsiakas, K.; Santer, R.; Nebert, D. W.; Rust, S.; Marquardt, T., SLC39A8 Deficiency: A Disorder of Manganese Transport and Glycosylation. Am J Hum Genet 2015, 97 (6), 894–903.

31. Jiang, Y.; Li, Z.; Sui, D.; Sharma, G.; Wang, T.; MacRenaris, K.; Takahashi, H.; Merz, K.; Hu, J., Rational engineering of an elevator-type metal transporter ZIP8 reveals a conditional selectivity filter critically involved in determining substrate specificity. Commun Biol 2023, 6 (1), 778.

32. Wiuf, A.; Steffen, J. H.; Becares, E. R.; Gronberg, C.; Mahato, D. R.; Rasmussen, S. G. F.; Andersson, M.; Croll, T.; Gotfryd, K.; Gourdon, P., The two-domain elevator-type mechanism of zinc-transporting ZIP proteins. Sci Adv 2022, 8 (28), eabn4331.

33. Zhang, Y.; Jiang, Y.; Gao, K.; Sui, D.; Yu, P.; Su, M.; Wei, G. W.; Hu, J., Structural insights into the elevator-type transport mechanism of a bacterial ZIP metal transporter. Nat Commun 2023, 14 (1), 385.

34. Zhang, Y.; Jafari, M.; Zhang, T.; Sui, D.; Sagresti, L.; Merz, K. M.; Hu, J., Molecular insights into substrate translocation in an elevator-type metal transporter. Nat Commun 2024, 15 (1), 9665.

35. Zhang, T.; Sui, D.; Zhang, C.; Cole, L.; Hu, J., Asymmetric functions of a binuclear metal center within the transport pathway of a human zinc transporter ZIP4. FASEB J 2020, 34 (1), 237–247.

36. Jiang, Y.; MacRenaris, K.; O’Halloran, T. V.; Hu, J., Determination of metal ion transport rate of human ZIP4 using stable zinc isotopes. J Biol Chem 2024, 300 (9), 107661.

37. Jeong, J.; Walker, J. M.; Wang, F.; Park, J. G.; Palmer, A. E.; Giunta, C.; Rohrbach, M.; Steinmann, B.; Eide, D. J., Promotion of vesicular zinc efflux by ZIP13 and its implications for spondylocheiro dysplastic Ehlers-Danlos syndrome. Proc Natl Acad Sci U S A 2012, 109 (51), E3530–8.

38. Zhao, M.; Zhou, B., A distinctive sequence motif in the fourth transmembrane domain confers ZIP13 iron function in Drosophila melanogaster. Biochim Biophys Acta Mol Cell Res 2020, 1867 (2), 118607.

39. Xiao, G.; Wan, Z.; Fan, Q.; Tang, X.; Zhou, B., The metal transporter ZIP13 supplies iron into the secretory pathway in Drosophila melanogaster. Elife 2014, 3, e03191.

40. Li, H.; Cui, Y.; Hu, Y.; Zhao, M.; Li, K.; Pang, X.; Sun, F.; Zhou, B., Mammalian SLC39A13 promotes ER/Golgi iron transport and iron homeostasis in multiple compartments. Nat Commun 2024, 15 (1), 10838.

41. Hu, J.; Jiang, Y., Evolution, classification, and mechanisms of transport, activity regulation, and substrate specificity of ZIP metal transporters. Crit Rev Biochem Mol Biol 2024, 59 (5), 245–266.

42. Polesel, M.; Ingles-Prieto, A.; Christodoulaki, E.; Ferrada, E.; Doucerain, C.; Altermatt, P.; Knecht, M.; Kuhn, M.; Steck, A. L.; Wilhelm, M.; Manolova, V., Functional characterization of SLC39 family members ZIP5 and ZIP10 in overexpressing HEK293 cells reveals selective copper transport activity. Biometals 2022, 36, 227–237.

43. Fersht, A., Structure and Mechanism in Protein Science: A Guide to Enzyme Catalysis and Protein Folding. W.H. Freeman and Company: New York, 2017; Vol. 9.

44. Arnold, F. H., Innovation by Evolution: Bringing New Chemistry to Life (Nobel Lecture). Angew Chem Int Ed Engl 2019, 58 (41), 14420–14426.

45. Yang, J.; Li, F.-Z.; Arnold, F. H., Opportunities and Challenges for Machine Learning-Assisted Enzyme Engineering. ACS Central Science 2024, 10 (2), 226–241.

46. Ji, C.; Kosman, D. J., Molecular mechanisms of non-transferrin-bound and transferring-bound iron uptake in primary hippocampal neurons. Journal of Neurochemistry 2015, 133 (5), 668–683.

47. Fisher, A. L.; Phillips, S.; Wang, C.-Y.; Paulo, J. A.; Xiao, X.; Moschetta, G. A.; Sridhar, A.; Mancias, J. D.; Babitt, J. L., Endothelial ZIP8 plays a minor role in BMP6 regulation by iron in mice. Blood 2024, 143 (23), 2433–2437.

48. Aydemir, T. B.; Liuzzi, J. P.; McClellan, S.; Cousins, R. J., Zinc transporter ZIP8 (SLC39A8) and zinc influence IFN-γ expression in activated human T cells. Journal of Leukocyte Biology 2009, 86 (2), 337–348.

49. Liu, M.-J.; Bao, S.; Gálvez-Peralta, M.; PyleCharlie J.; RudawskyAndrew C.; PavloviczRyan E.; KillileaDavid W.; Li, C.; NebertDaniel W.; WewersMark D.; KnoellDaren L., ZIP8 Regulates Host Defense through Zinc-Mediated Inhibition of NF-κB. Cell Reports 2013, 3 (2), 386–400.

50. Steimle, B. L.; Smith, F. M.; Kosman, D. J., The solute carriers ZIP8 and ZIP14 regulate manganese accumulation in brain microvascular endothelial cells and control brain manganese levels. Journal of Biological Chemistry 2019, 294 (50), 19197–19208.

51. Lin, W.; Vann, D. R.; Doulias, P.-T.; Wang, T.; Landesberg, G.; Li, X.; Ricciotti, E.; Scalia, R.; He, M.; Hand, N. J.; Rader, D. J., Hepatic metal ion transporter ZIP8 regulates manganese homeostasis and manganese-dependent enzyme activity. Journal of Clinical Investigation 2017, 127 (6), 2407–2417.

