## Supplementary Information for "Targeting the selectivity filter to drastically alter the activity and substrate spectrum of a promiscuous metal transporter"

*Equally contributive to this work

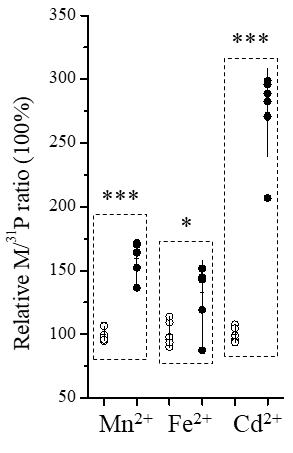

**Figure S1.** Detection of the activities for Mn^2+^, Fe^2+^, and Cd^2+^ under the optimized condition. For each metal, the relative M/^31^P ratio of the ZIP8 group (solid circle) is expressed as a percentage of the M/^31^P ratio of the empty vector group (open circle). Each data point represents the result of one sample, and six replicates were conducted under one condition. For the result of each metal, the horizontal bar is the mean of three replicates and the vertical bar shows 1±S.D. Two tailed student’s *t* test was used to test statistical significance. *: *P*<0.05; ***: *P*<0.001. The *P* values for Mn^2+^, Fe^2+^, and Cd^2+^, are 1x10^-6^, 0.01, and 2x10^-7^, respectively.

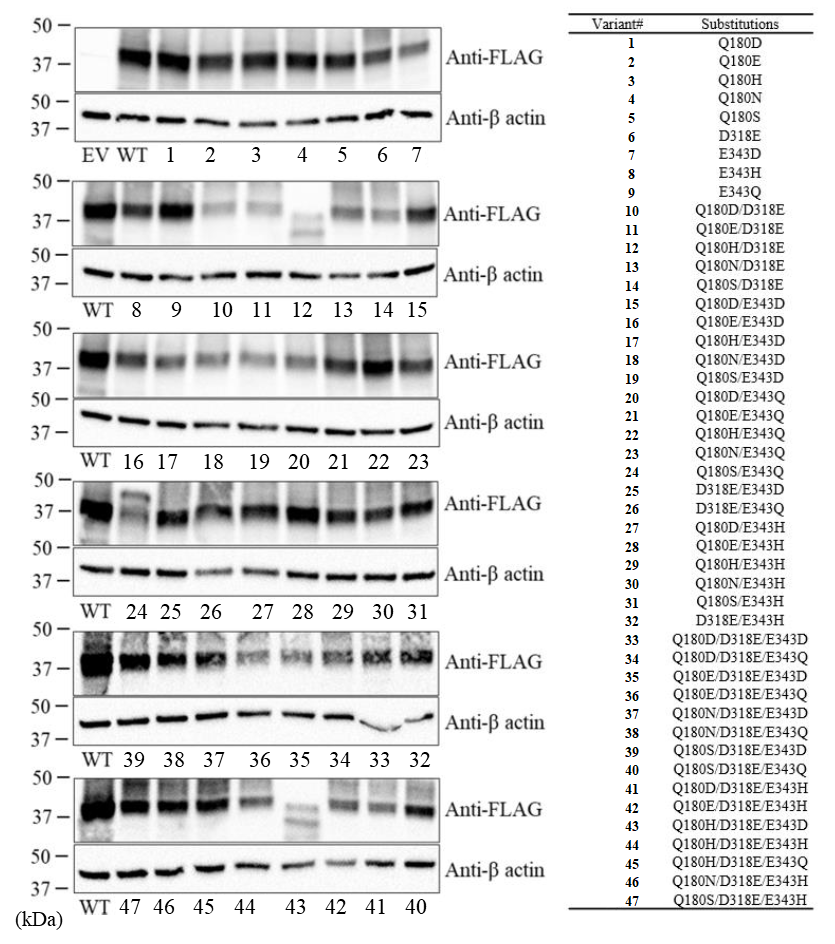

**Figure S2.** Expression of N-terminal FLAG-tagged ZIP8 and its variants detected by Western blot. An anti-FLAG antibody was used to detect the total expression of ZIP8 and its variants. β-actin was used as the loading control. Variants #12 (Q180H/D318E), #24 (Q180S/E343Q), and #43 (Q180H/D318E/E343D) showed different molecular weights from other constructs and exhibited poor activities (**Table 1**).

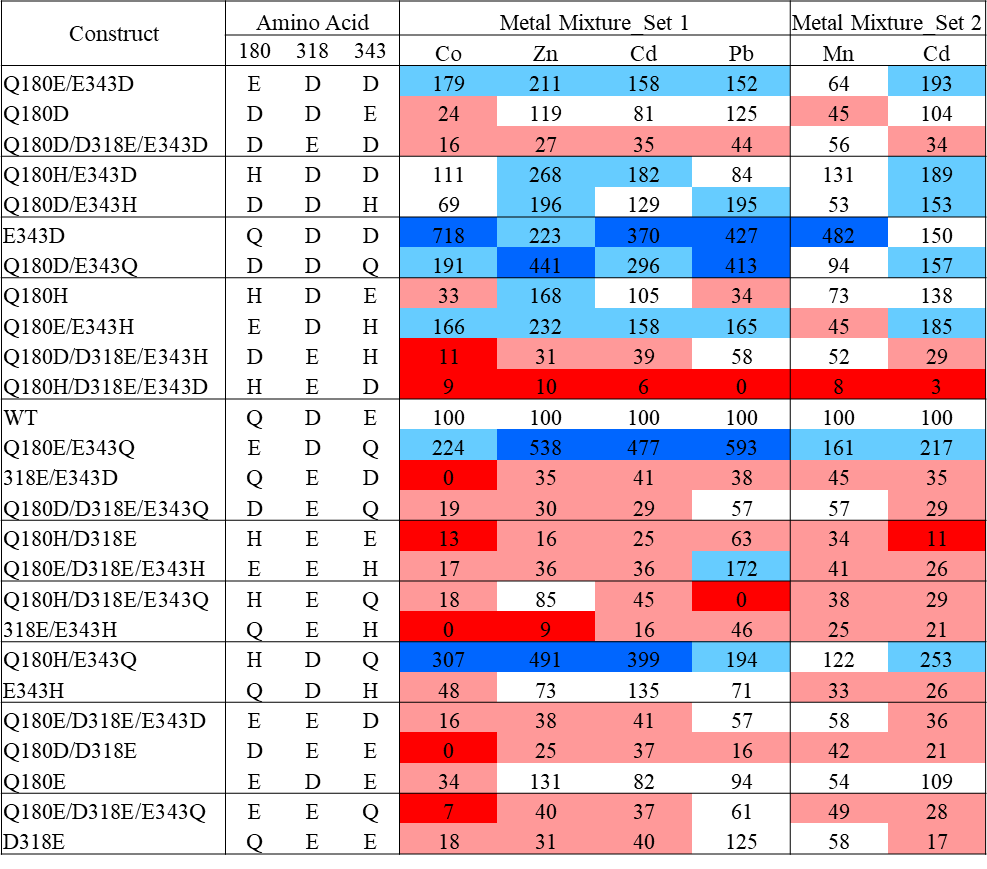

**Figure S3.** Comparison of the constructs with the same amino acid composition at the selectivity filter. The constructs in the same frame have the same amino acid composition but different distribution at the positions of 180, 318, and 343. They often do not exhibit the same activity or substrate preference, suggestive of non-overlapping roles of these residues in transport. Data are retrieved from **Table 1**.

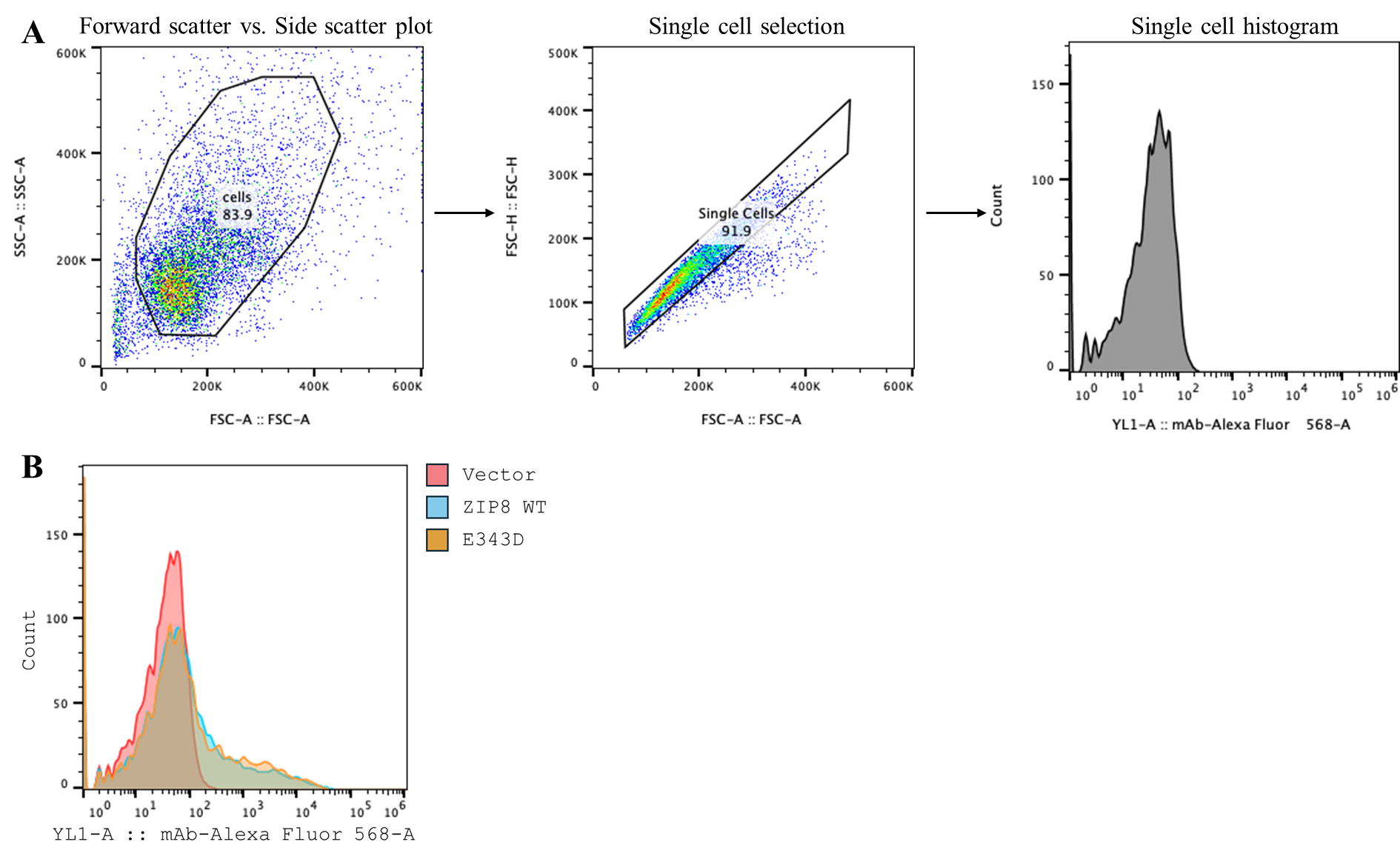

**Figure S4.** Flow cytometry data processing. (**A**) Representative data processing. (**B**) Comparison of cell surface expression levels of empty vector, ZIP8, and the E343D variant in a histogram chart.
